# MiR-203 inhibits the proliferation, invasion, and migration of pancreatic cancer cells by down-regulating fibroblast growth factor 2

**DOI:** 10.1101/2020.04.06.027326

**Authors:** Xi-Feng Fu, Hai-Chao Zhao, Chang-Zhou Chen, Kang Wang, Fei Gao, Yang-Zhang Tian, Hui-Yu Li, Hao-Liang Zhao

## Abstract

**Background:** Aberrant fibroblast growth factor 2 (FGF2) expression is a major cause of poor prognosis in pancreatic cancer. MiR-203 is a newly discovered microRNA (miRNA) that can affect the biological behavior of tumors. This study investigated whether miR-203 can regulate FGF2 expression and its role in pancreatic cancer cell proliferation, apoptosis, invasion, and migration.

**Methods:** MiR-203 expression in different cell lines was examined by qRT-PCR, followed by the establishment of knockdown and overexpression cell models. We used the CCK-8 assay to examine cell proliferation and the annexin V-APC/7-AAD double-staining method to detect apoptosis. In addition, we used wound healing and transwell assays to investigate the effects of miR-203 on the migration and invasion of pancreatic cancer cells. The effects of miR-203 knockdown and overexpression on FGF2 mRNA expression were detected by qRT-PCR. We also overexpressed FGF2 and examined the effects of FGF2 overexpression on the proliferation, apoptosis, invasion, and migration of pancreatic cancer cells. The binding of miR-203 to FGF2 was assessed by a luciferase reporter assay.

**Results:** We found that the miR-203 expression level was significantly down-regulated in pancreatic cancer cells compared to normal pancreatic cells. Functionally, the knockdown of miR-203 inhibited cell proliferation and increased apoptosis. Equally important, miR-203 reduced the migration and invasion of pancreatic cancer cells. In addition, we found that miR-203 overexpression inhibited FGF2 expression in pancreatic cancer cells by qRT-PCR. FGF2 overexpression significantly affected the proliferation, invasion, and metastasis of pancreatic cancer cells. Mechanistically, miR-203 base-paired with the FGF2 mRNA, resulting in the knockdown of the FGF2 mRNA and the down-regulation of the FGF2 protein.

**Conclusions:** MiR-203 inhibits FGF2 expression, regulates the proliferation of pancreatic cancer cells, and inhibits the invasion and metastasis of pancreatic cancer cells.

## Introduction

Pancreatic cancer is currently a major disease that endangers human health, and its incidence is rapidly rising. Aggressive metastasis is the most important factor in the high mortality rate of patients with pancreatic cancer. However, the mechanism of metastasis is still unknown. The target molecules that underline metastasis are of great significance in the treatment of pancreatic cancer. Pancreatic ductal adenocarcinoma (PDAC) is the most common type of pancreatic cancer and one of the most challenging malignant tumors to treat. The median survival time of PDAC after diagnosis is 2–8 months, and the 5-year overall survival rate is less than 7% ^[1,2]^. The poor prognosis of pancreatic cancer is mainly due to the aggressiveness of cancer cells, early metastasis, and non-responsiveness to most chemotherapy regimens ^[3]^. Presently, surgical resection is the only feasible treatment for PDAC. However, less than 20% of tumors are resectable at the time of diagnosis ^[4]^. Furthermore, patients who undergo surgery may relapse, and the average survival time of patients undergoing resection is 12–20 months ^[5]^. Most pancreatic cancer patients are identified with early or locally advanced metastasis at the time of diagnosis, and the only effective treatment for these patients is chemoradiation. However, pancreatic cancer cells respond poorly to both chemotherapy and radiation ^[6,7]^. Although gemcitabine-based chemotherapy, as the standard treatment for advanced pancreatic cancer, can improve the prognosis, its effectiveness is limited because cancer cells often become drug resistant. Therefore, further pancreatic cancer research is urgently needed, and new diagnostic and therapeutic approaches are required to improve the prognosis of this disease.

Non-coding RNAs (ncRNAs) play extremely important roles in tumor development. MicroRNAs (miRNAs) are highly conserved single-stranded ncRNA molecules with a length of 18–25 nucleotides. MiRNAs can regulate gene expression by base-pairing with 3’ untranslated regions (3’ UTR), thereby enhancing mRNA degradation or inhibiting post-transcriptional translation ^[8]^. More than 2,500 miRNAs have been identified in plants, animals, and viruses. After miRNAs are produced in the nucleus, they are delivered into the cytoplasm by nuclear transporters and then guided into the RNA-induced silencing complex (RISC) where they facilitate target gene mRNA degradation or inhibit translation through complementary pairing with target gene mRNA bases. As more miRNAs are discovered, investigators are realizing that miRNAs play important roles in tumor development ^[9,10]^. For example, recent studies have reported the involvement of various miRNAs (e.g., miR-21, miR-155, and miR-210) in the development and progression of pancreatic cancer ^[11,12]^.

MiR-203 is located on chromosome 14*q*32.33. Compared to normal tissues, miR-203 shows abnormal expression in diverse malignancies, including bladder cancer, non-small-cell lung cancer, and endometrial cancer ^[13-15]^. Many studies have demonstrated that miR-203 plays important roles in tumor cell growth, migration, and invasion ^[16,17]^. For example, miR-203 is overexpressed in pancreatic cancer and not in normal pancreatic tissue and chronic pancreatitis, suggesting that miR-203 may be related to specific characteristics of tumors and their behavior ^[18]^. Other studies have suggested that FGF2 expression may be affected by miR-203 ^[19,20]^. However, the relevant mechanisms of action and signaling pathways in pancreatic cancer remain unknown. In this study, we established cell models of miR-203 knockdown and overexpression to explore the regulatory effects of miR-203 on the expression of FGF2 as well as the proliferation, invasion, and migration of pancreatic cancer cells.

## Patients and Methods

### Key reagents and materials

Pancreatic cancer cell lines (PANC-1, AsPC-1, BxPC-3, and HPAC) were purchased from the Shanxi Academy of Medical Sciences (Taiyuan, China). Normal human pancreatic epithelial cells (HPDE 6-C7 and HEK293T) were purchased from the Chinese Academy of Sciences (Shanghai, China). Dulbecco’s Modified Eagle’s Medium (DMEM), fetal bovine serum (FBS), and Opti-MEM medium were procured from Gibco (Grand Island, NY, USA). Lipofectamine 2000 was procured from Invitrogen (Carlsbad, CA, USA). The PrimeScript^™^ RT Kit and SYBR Green dye were procured from TaKaRa (Otsu, Shiga, Japan). Mimic-miR-203-3p, mimic-NC, inhibitor-miR-203-3p, and inhibitor-NC were obtained from RiboBio (Guangzhou, China). The pGL3 vector and Dual-Luciferase Activity Assay were purchased from Promega (Madison, WI, USA). The EdU-Alexa Fluor^®^ 488 Cell Proliferation Assay was purchased from Thermo Fisher Scientific (Waltham, MA, USA). The Annexin V-Fluorescein Isothiocyanate (FITC)/Propidium Iodide (PI) Double-Staining Cell Apoptosis Assay was purchased from Nanjing Kaiji (Nanjing, China).

### Cell culture and cell transfection

PANC-1, AsPC-1, BxPC-3, HPAC, and HPDE cells were cultured in DMEM containing 10% FBS and 1% streptomycin at 37°C in an atmosphere of 5% CO_2_. The cells were passaged at 1:4 or 1:5, and then used for experiments at the logarithmic growth phase. The cells were digested with 0.25% trypsin. The digested cells were counted and diluted to a concentration of 5 × 10^4^ cells/mL. Thereafter, 100 μL of the suspension was added into each well of a 96-well culture plate, which was then incubated at 37°C in an atmosphere of 5% CO_2_ for 24 h. For cell transfection, 0.25 μg of siRNA was diluted with 25 μL of serum-free Opti-MEM, mixed gently, and incubated at room temperature for 5 min, and 0.5 μL of Lipofectamine 2000 was diluted with 25 μL of serum-free Opti-MEM, mixed gently, and incubated at room temperature for 5 min. Both components were subsequently combined, mixed gently, and incubated for 20 min at room temperature. The siRNA–Lipofectamine 2000 mixture was then added into the wells, which contained 50 μL of medium, and gently mixed. After 4–6 h of incubation, the media containing the siRNA–Lipofectamine 2000 mixture was carefully aspirated and replaced with fresh media. The culture plate was incubated as indicated above for 24 h. Thereafter, the cells were treated with mimic-NC, mimic-miR203a-3p, inhibitor-NC or inhibitor-miR203a-3p.

### Detection of cell proliferation by the CCK-8 assay

The cells were digested, counted, and diluted to a concentration of 5 × 10^4^ cells/mL. Thereafter, 100 μL of the cell suspension was added into each well of a 96-well cell culture plate, which was then incubated at 37°C in an atmosphere of 5% CO_2_ for 24 h. For cell transfection, 0.25 μg of miRNA was diluted with 25 μL of serum-free Opti-MEM, mixed gently, and incubated at room temperature for 5 min, and 0.5 μL of Lipofectamine 2000 was diluted with 25 μL of serum-free Opti-MEM, mixed gently, and incubated at room temperature for 5 min. Both components were subsequently combined, mixed gently, and incubated at room temperature for 20 min. The miRNA– Lipofectamine 2000 mixture was then added into the wells, which contained 50 μL of medium, and mixed gently. After 4–6 h of incubation, the media containing the miRNA–Lipofectamine 2000 mixture was carefully aspirated and replaced with fresh media. The culture plate was incubated as indicated above for 24 h. Thereafter, 10 μL of CCK-8 reagent was added into each well. The culture plate was incubated for 3 h and mixed gently on a shaker for 10 min. A microplate reader was used to read the OD value of each well at a λ of 450 nm. The inhibition rate (%) was calculated as follows: (negative control group - experimental group) / negative control group × 100%.

### Detection of apoptosis by the annexin V-APC/7-AAD double-staining method

The cells were washed twice with phosphate-buffered saline (PBS), trypsinized, and centrifuged at 1000 rpm for 5 min to collect 5 × 10^5^ cells, which were then resuspended in 500 μL of binding buffer. Thereafter, 5 μL of annexin V-APC was added to the cell suspension, followed by mixing, and then 5 μL of 7-AAD was added, again followed by mixing. The samples were incubated at room temperature in the dark for 5–15 min, and apoptosis was detected by flow cytometry.

### Detection of migration and invasion by cell wound healing and transwell assays

For cell wound healing assays, cells at the logarithmic growth phase were cultured to confluence in 6-well plates. The next day, when the cell concentration reached approximately 60%, the cell layer was scratched with a 200-μL micropipette tip in the center of the well. The cells were gently rinsed with PBS and incubated with 1% FBS-containing medium. After 24 h of incubation, the cells were removed from the incubator, photographed (magnification, 200×), and the cell migration distance was measured. The width of the wound healing site was quantified and compared with baseline values. All experiments were repeated independently in triplicate.

For transwell assays, cells were removed from serum and starved for 24 h using incomplete medium. Matrigel was thawed at 4°C overnight and 2 × 10^4^ cells from each group in 200 μL of serum-free medium were seeded in the upper chamber (pore size, μm; Corning, NY, USA) without (migration) or with (invasion) Matrigel (BD Biosciences, San Jose, CA, USA). Thereafter, 600 μL of RPMI-1640 medium containing 10% FBS was added to the lower chamber. After 24 h of incubation, the upper chambers were fixed with 4% polymethanol for 30 min, and then stained with 0.1% crystal violet for 30 min. The cells that migrated through the membrane and invaded the underside of the upper chamber were photographed. Five random fields were selected to calculate the number of cells that underwent migration or invasion.

### qRT-PCR

Total RNA was extracted using TRIzol reagent (Invitrogen) and reversely transcripted into cDNA using the PrimeScript^™^ RT Kit. qRT-PCR PCR was performed on the Bio-Rad CFX96 System (Hercules, CA, USA). The reactions consisted of 5.0 μL of 2× SYBR Green Master Mix, 0.5 μL each of forward and reverse primers (2.5 μM), 1 μL of cDNA, and RNase-free double-distilled H_2_O with reaction conditions as follows: 95°C for 5 min, followed by 40 cycles at 95°C for 15 s and 60°C for 1 min. The relative expression of target mRNAs was quantified using the 2^-ΔΔCt^ method.

### Dual-luciferase reporter assay

Using the HEK293T cell genome as the template, the PCR product of the FGF2 3’-UTR full-length fragment or mutant fragment was double-digested, and then ligated into the pGL3 vector. The luciferase reporter plasmids were cotransfected into cells with pGL3-FGF2-WT (or pGL3-FGF2-MUT) together with miR-203 mimic (or miR-NC) by using Lipofectamine(tm) 2000 reagent. After 36 h of incubation, firefly luciferase and renilla luciferase activities were detected according to the instructions of the Dual-Glo Luciferase Assay System Kit. All experiments were repeated independently in triplicate.

### Statistical analysis

SPSS 26.0 Software (IBM, Armonk, NY, USA) was used for statistical analysis. The data were analyzed by the χ^2^ test or Fisher’s exact test. The correlations were analyzed by Pearson’s test (r, P). Paired and unpaired continuous variables were compared by the Student’s *t*-test or Mann–Whitney U test. *P*-values < 0.05 were considered statistically significant in all tests. All graphs were produced using GraphPad Prism (GraphPad Software Inc., San Diego, CA, USA) and ImageJ 2X (Rawak Software Inc., Stuttgart, Germany).

## Results

### miRNA-203 expression in different cell lines

We used qRT-PCR to quantify miRNA-203-3p expression in PANC-1, AsPC-1, BxPC-3, HPAC, and HPDE cell lines and found that the miR-203 level was lower in pancreatic cancer cells than that in HPDE cells. Furthermore, miRNA-203-3p expression was the lowest in AsPC-1 cells (**Fig. 1a**), indicating that the AsPC-1 cell line was suitable for further experiments.

**Fig. 1.**
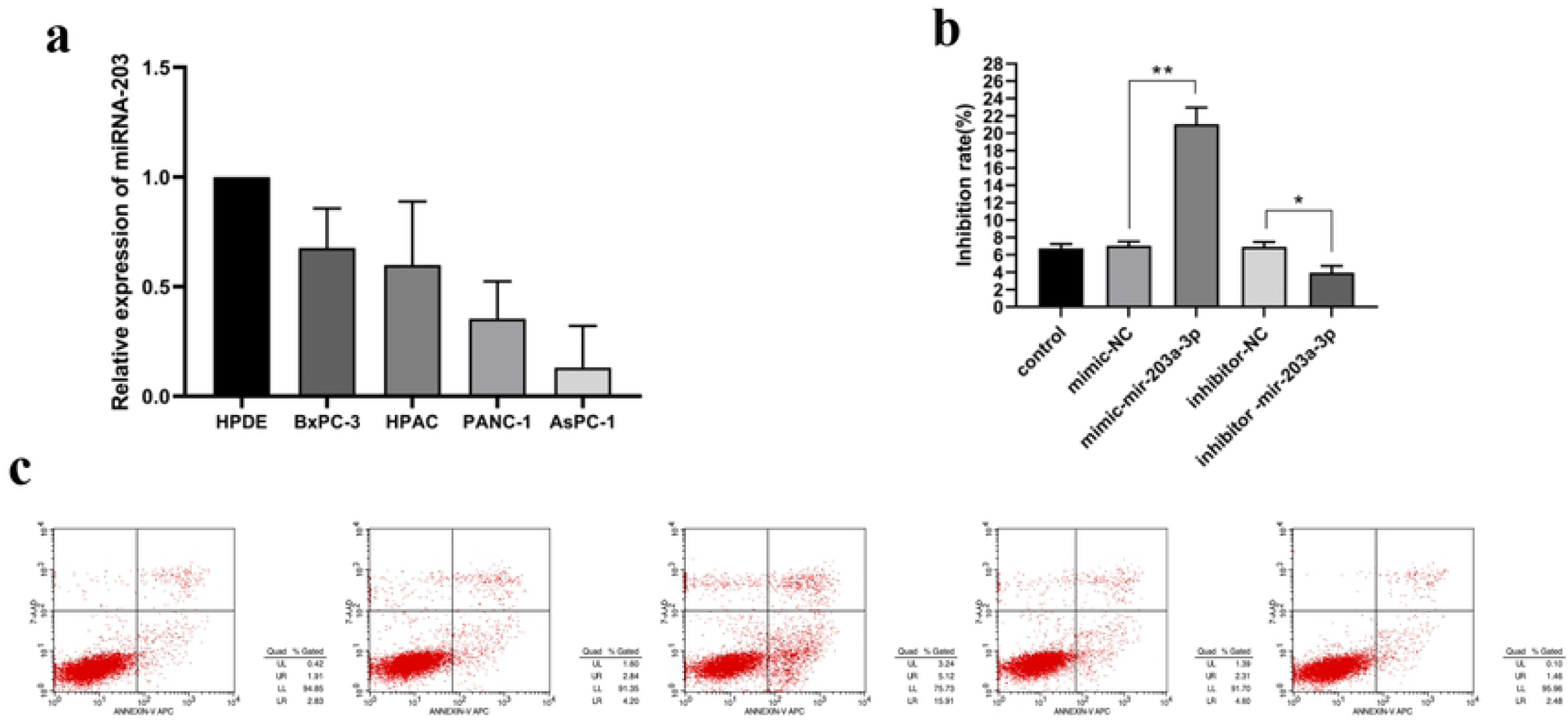
miR-203a-3p overexpression increases the apoptosis of pancreatic cancer cells. **a** Relative expression of miRNA-203a-3p in pancreatic cancer cell lines by qRT-PCR. **b** and **c** miR-203a-3p overexpression increases pancreatic cancer cell apoptosis as detected by the annexin V-APC/7-AAD double-staining method. All data are presented as the mean ± SEM of three experiments. \**p* < 0.05, \*\**p* < 0.01.

### miR-203 inhibits the proliferation of AsPC-1 cells

To investigate the role of miRNA-203-3p in pancreatic cancer cells, we used inhibitor-miR-203-3p, which targeted the junction sites of miR-203-3p. We also generated overexpression plasmids of miR-203-3p for transfection into AsPC-1 cells. The results of the CCK-8 assay showed that miR-203-3p could inhibit the proliferation of AsPC-1 cells. As shown in **Table 2**, compared with the mimic-NC group, the inhibition rate of the mimic-miR203-3p group increased significantly. However, compared with the inhibitor-NC group, the inhibition rate of the inhibitor-miR203-3p group decreased significantly.

### miR-203 increases apoptosis of pancreatic cancer cells

The AsPC-1 cell apoptosis rate was not significantly different among control, mimic-NC, and inhibitor-NC groups. Compared with the mimic-NC group, the apoptosis rate of the mimic-miR-203-3p group increased significantly. However, compared with the inhibitor-NC group, the apoptosis rate of the inhibitor-miR-203-3p group was decreased slightly (**Fig. 1b and c, Table 3**).

### miR-203-3p reduces the invasion and migration of pancreatic cancer cells

We examined the ability of different types of cells to invade and migrate. Compared with the mimic-NC group, the mimic-miR203a-3p group showed increased migration and invasion at 24 h. Nevertheless, overexpression of miR203a-3p promoted the migration and invasion of AsPC-1 cells. These results were opposite to those of the inhibitor-miR203a-3p group (**Tables 4 and 5, Fig. 2a, b, c** and **d**). Taken together, these findings suggest that miR203a-3p promotes the progression of gastric cancer *in vivo*.

**Fig. 2.**
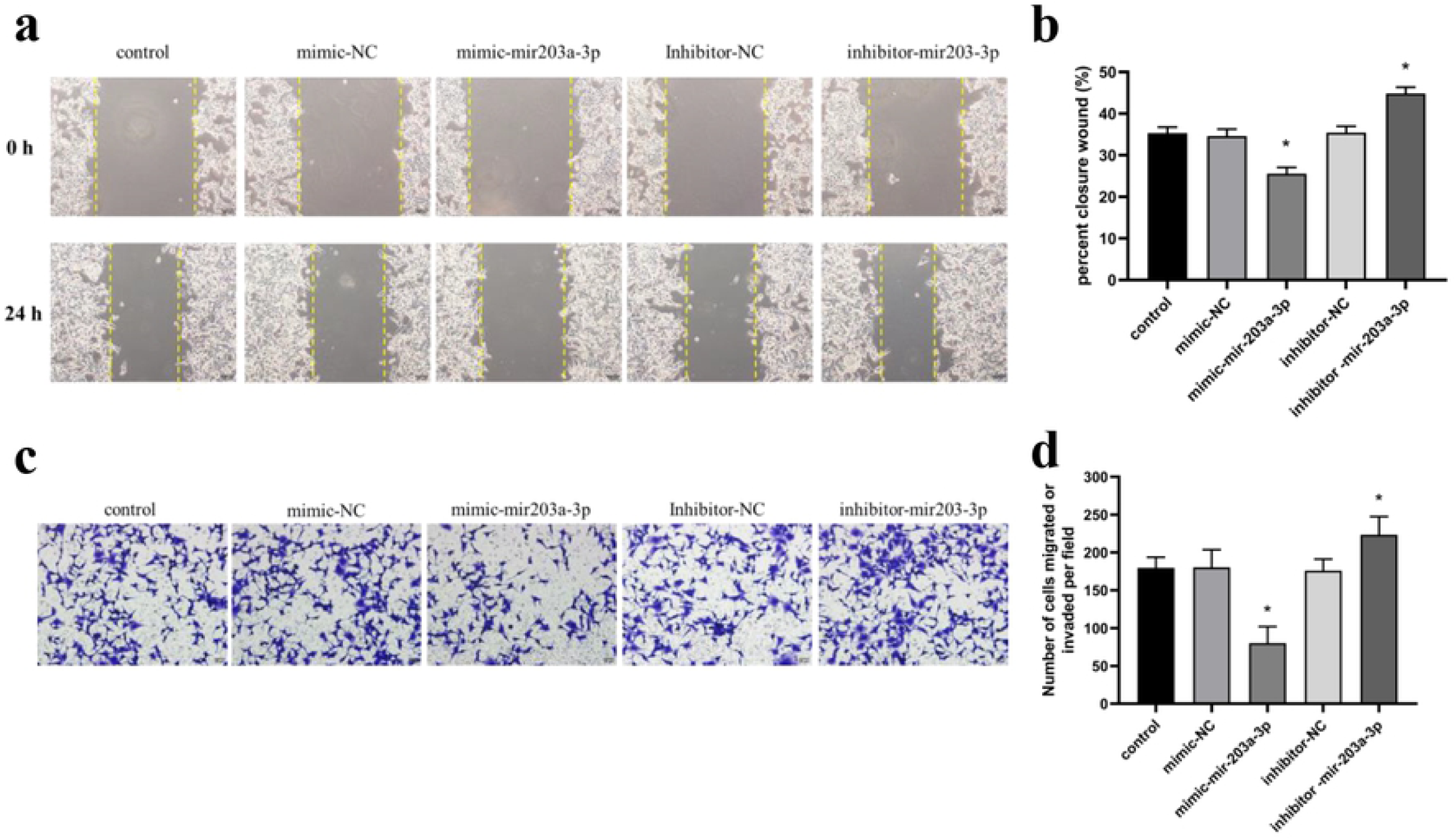
miR-203a-3p overexpression reduces the migration and invasion of pancreatic cancer cell lines. **a** and **b** The effects of miR-203a-3p overexpression on migration were detected by wound healing assays. **c** and **d** The effects of miR-203a-3p overexpression on invasion were detected by transwell assays. All data are presented as the mean ± SEM of three experiments. \**p* < 0.05.

### miRNA-203 inhibits FGF2 expression in pancreatic cancer cell lines

miRNA-203 expression was reduced significantly by inhibitor-miR-203a-3p, whereas miRNA-203 expression was enhanced by mimic-miR-203a-3p. We found that miRNA-203 overexpression in AsPC-1 cells decreased the FGF2 mRNA level. By contrast, miRNA-203 knockdown in AsPC-1 cells increased the FGF2 mRNA level, as determined by qRT-PCR. These results indicate that up-regulated miRNA-203 elicited inhibitory effects on FGF2 expression (**Table 6, Fig. 3a**).

**Fig. 3.**
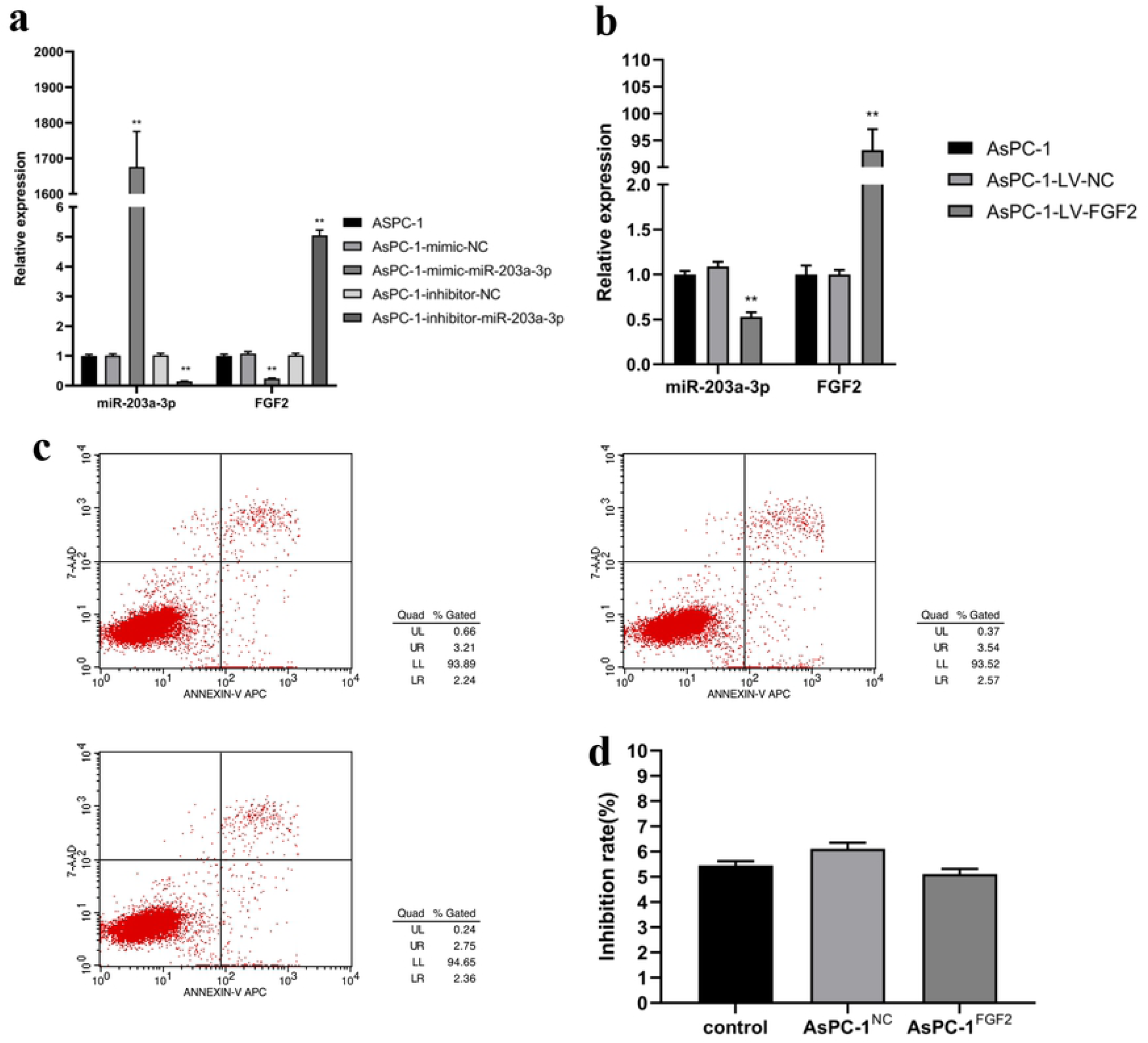
miRNA-203a-3p overexpression inhibits FGF2 expression in pancreatic cancer cell lines. **a** Relative expression of miRNA-203-3p and FGF2 in pancreatic cancer cell lines after miRNA-203a-3p overexpression and knockdown by qRT-PCR. **b** Relative expression of miRNA-203a-3p and FGF2 in pancreatic cancer cell lines after FGF2 overexpression and knockdown by qRT-PCR. **c** and **d** FGF2 induced apoptosis of pancreatic cancer cells as detected by the annexin V-APC/7-AAD double-staining method. All data are presented as the mean ± SEM of three experiments. \**p* < 0.05, \*\**p* < 0.01.

To verify the role of FGF2 in AsPC-1 cells, we simultaneously transfected the control plasmid and FGF2 overexpression plasmid and performed qRT-PCR. We found that the FGF2 mRNA level in the overexpression group increased significantly (**Table 7, Fig. 3b**).

### FGF2 significantly affects the proliferation of AsPC-1 cells

FGF2 expression in pancreatic cancer cell lines significantly affects cell proliferation. Compared with AsPC-1 and AsPC-1NC groups, the inhibition rate of the AsPC-1FGF2 group was −15.68%, indicating that FGF2 expression can promote the proliferation of AsPC-1 cells (**Table 8**).

### FGF2 affects AsPC-1 cell apoptosis

The results of flow cytometry experiments showed that there was no significant difference in the apoptosis rate among AsPC-1, AsPC-1^NC^, and AsPC-1^FGF2^ groups, indicating that the change in FGF2 expression did not affect apoptosis (**Fig. 3c** and **d**, **Table 9**).

### FGF2 overexpression increases the migration and invasion of pancreatic cancer cells

Compared with control and AsPC-1^NC^ groups, the wound closure ratio, an indicator of cell migration, of the AsPC-1^FGF2^ group increased (**Fig. 4a** and **b**, **Table 10**). Compared with control and AsPC-1^NC^ groups, the invasion cell number of the AsPC-1^FGF2^ group increased significantly (**Fig. 4c** and **d**, **Table 11**). These results indicate that FGF2 can promote the ability of the AsPC-1 pancreatic cancer cell line to migrate and invade.

**Fig. 4.**
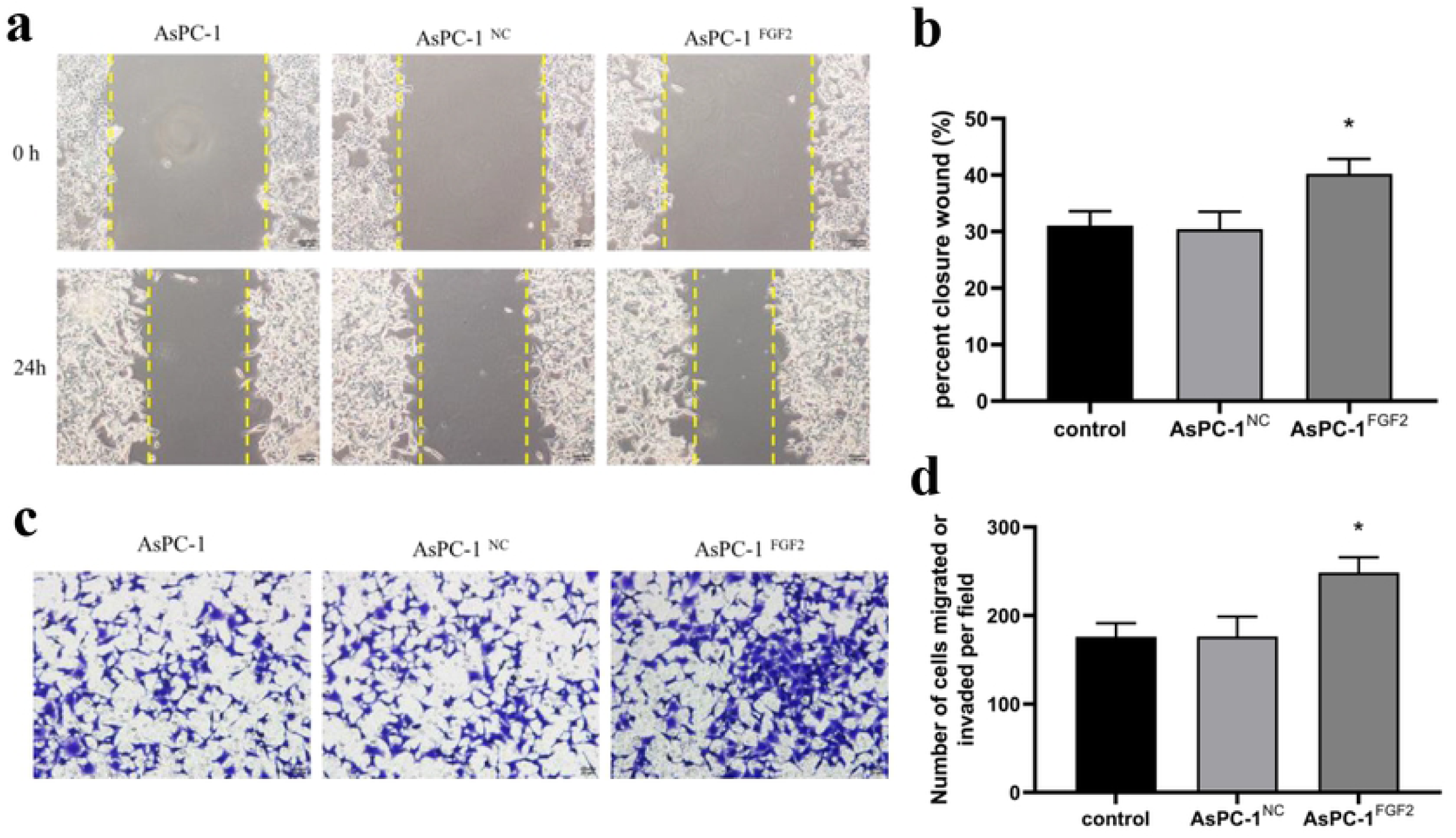
FGF2 overexpression? increases the migration and invasion of pancreatic cancer cell lines. **a** and **b** The effects of FGF2 overexpression on migration were detected by wound healing assays. **c** and **d** The effects of FGF2 overexpression on invasion were detected by transwell assays. All data are presented as the mean ± SEM of three experiments. \**p* < 0.05.

### miR-203a-3p directly targets FGF2 in pancreatic cancer cells

According to a bioinformatics query (TargetScan), the predicted site of action was found to be at the site of miR-203a-3p binding to the 3’ UTR region of FGF2. We used a dual luciferase reporter system to verify whether miR-203a-3p interacts with the seed region of the 3’ UTR region of FGF2 (**Fig. 5a**). We successfully constructed a dual luciferase reporter gene vector for the target gene seed region (including wild-type and mutant) and co-transfected it with miR-203a-3p overexpression and negative control vectors. The cell lines were tested for luciferase activity in transfected cells. The results showed that luciferase activity in AsPC-1 cells co-transfected with miR-203-3p overexpression and wild-type vectors was significantly reduced compared to luciferase activity in control-transfected cells. These results indicate that miR-203-3p interacts directly with the 3’ UTR region of FGF2. This mechanism of direct binding indicates that miR-203-3p directly regulates the target gene FGF2 (**Fig. 5b**).

**Fig. 5.**
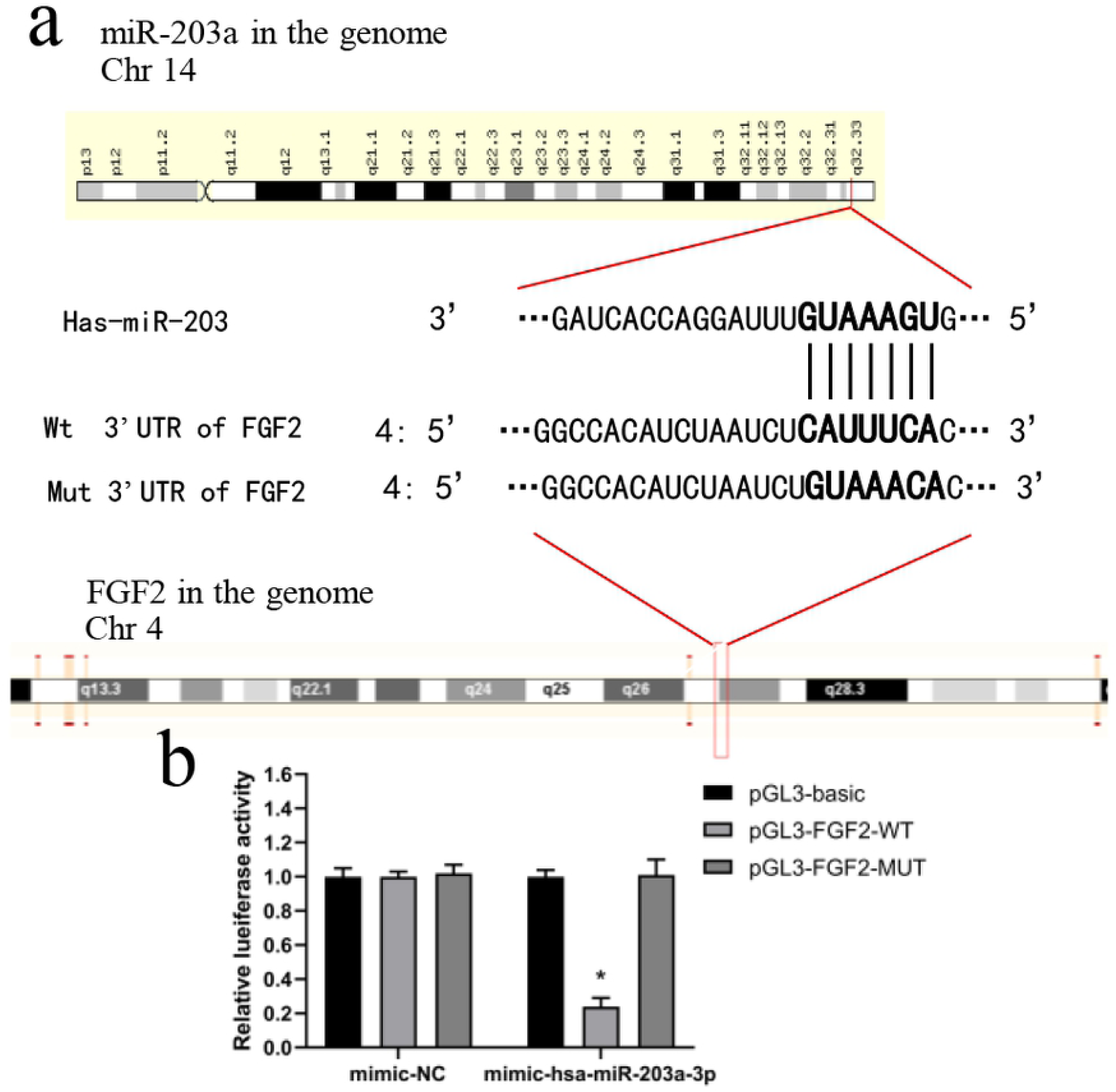
miR-203 directly targets FGF2 in pancreatic cancer cells. **a** Sequence alignment of miR-203a-3p and 3’ UTR of FGF2. **B** The resuts of the luciferase assay co-transfected with miR-203 mimic and a luciferase reporter containing the FGF2 3’ UTR (Wt) or a mutant (Mut). All data are presented as the mean ± SEM of three experiments. \**p* < 0.05.

## Discussion

Fibroblast growth factor 2 (FGF2), also known as basic fibroblast growth factor (bFGF) and FGF-β, is a growth factor and signaling protein encoded by the FGF2 gene ^[21]^. The protein encoded by this gene is a member of the fibroblast growth factor (FGF) family. FGF family members bind heparin and possess broad mitogenic and angiogenic activities ^[22]^. FGF2 has also been implicated in diverse biological processes, such as limb and nervous system development, wound healing, and tumor growth ^[23]^. The mRNA for this gene contains multiple polyadenylation sites and is alternatively translated from non-AUG (CUG) and AUG initiation codons, resulting in five different isoforms with distinct functions. The CUG-initiated isoforms are localized in the nucleus and responsible for the intracrine effect of FGF, whereas the AUG-initiated form is mostly cytosolic and responsible for paracrine and autocrine effects ^[24]^. FGF2 binds to specific cell surface tyrosine kinase receptors (FGFR) to activate intracellular signaling, thereby affecting the proliferation, migration, or differentiation of many types of cells. Kostas et al. ^[25]^ described the anti-apoptotic effects of FGF1 and FGF2 in cells, which was independent of FGFR activation and downstream signaling. Several investigators showed that FGF2 can promote epithelial-mesenchymal transition (EMT) in injured tissues ^[26]^. We found that the heparanase/syndecan-1 (HPA/SDC1) axis can up-regulate the FGF2 level and increase the expression of downstream palladin by activating the phosphoinositide 3-kinase/Akt (PI3K/Akt) signaling pathway, thereby leading to the activation of EMT. We also reported that EMT promotes the migration and invasion of pancreatic cancer cells ^[27]^.

MicroRNAs (miRNAs) are novel, highly stable, and abundant endogenous noncoding RNAs. With the development of high-throughput sequencing and bioinformatics analysis, an increasing number of miRNAs has been identified and confirmed to regulate the development and progression of various human cancers in recent years ^[28,29]^. However, few miRNAs have been functionally and mechanistically characterized with regard to gastric cancer, and the biological functions of most miRNAs have yet to be explored. In this study, we identified a novel metastasis-related miRNA, miR-203, which was significantly down-regulated in pancreatic cancer cells. We overexpressed miR-203 in AsPC-1 cells by transfection and observed that the proliferation and apoptosis of pancreatic cancer cells were decreased. Equally important, miR-203 overexpression could inhibit the invasion and metastasis of AsPC-1 cells.

We previously demonstrated that FGF2 overexpression could contribute to the resistance to chemotherapy of pancreatic cancer cells ^[30,31]^. Patients with pancreatic cancer and high FGF2 expression are not responsive to postoperative chemotherapy with gemcitabine, and overall, the prognosis is poor ^[32]^. In this study, we found that increased miR-203 expression could inhibit the transcription of the FGF2 mRNA in AsPC-1 cells, thereby improving the prognosis of patients with pancreatic cancer. In recent years, the relationship between cytokines and tumorigenesis has gained momentum. Cytokine–receptor interactions activate various signaling pathways in cells and play important roles in regulating cell proliferation, apoptosis, and angiogenesis ^[33]^. FGF2, a growth-promoting factor, has been found to be highly expressed in breast cancer, gastric cancer, and thyroid cancer, as well as other normal tissues, and aberrant FGF2/FGF2R signaling may promote tumor development ^[34]^. MiRNAs regulate gene expression after transcription. Therefore, miRNAs are key regulators of gene expression and promising candidates for biomarker development. MiRNAs also guide the RISC to degrade mRNAs or prevent translation by base-pairing with target gene mRNAs ^[8]^. In this study, we found that miR-203-3p, combined with the 3’ UTR region of FGF2, could exert a direct regulatory effect. These results are consistent with those of previous studies ^[35, 36]^, and they explain why miR-203 can affect the proliferation, invasion, and migration of pancreatic cancer cells. However, we also found that down-regulation of miR-203 inhibited AsPC-1 cell apoptosis, but overexpression of FGF2 had no significant effect on cell viability. It is possible that miR-203 affects the viability of AsPC-1 cells through other pathways, and further studies are needed to explore this hypothesis.

Functional studies have confirmed that miRNA dysregulation is causal in many cases of cancer, with miRNAs acting as tumor suppressors or oncogenes (onco-miRs) ^[37]^, and miRNA mimics and molecules that target miRNAs (anti-miRs) showing promise in preclinical development. Several miRNA-targeted therapeutics have reached clinical development, including a mimic of the tumor suppressor miRNA miR-34, which reached phase I clinical trials for treating cancer, and anti-miRs targeted at miR-122, which reached phase II trials for treating hepatitis ^[38, 39]^.

There is still room for further studies. CircRNAs have been reported to have multiple and diverse molecular mechanisms in the development and progression of various cancers, among which ceRNA is the most important. The ceRNA hypothesis states that circRNAs serve as ceRNAs to positively regulate the expression of miRNA target genes ^[40, 41]^. For example, circAKT3 promotes PIK3R1 expression via miR-198 suppression in gastric cancer ^[42]^. CircPSMC3 acts as a miR-296-5p sponge to promote the expression of phosphatase and tensin homolog ^[43]^. In subsequent studies, we intend to use miR-203 as a starting point to identify circRNAs that can affect miR-203, so as to further explore the targets of pancreatic cancer treatment.

## Acknowlegment

We would like to thank TopEdit (www.topeditsci.com) for English language editing of this manuscript.

